# GABA_A_ α Subunit Control of Hyperactive Behavior in Developing Zebrafish

**DOI:** 10.1101/2021.07.01.450600

**Authors:** Wayne Barnaby, Hanna E. Dorman Barclay, Akanksha Nagarkar, Matthew Perkins, Gregory Teicher, Josef G. Trapani, Gerald B. Downes

**Author notes:** Author for correspondence: Gerald B. Downes, 611 North Pleasant St., Morrill Science Center, Building 4 North, Amherst MA 01002.

## Abstract

GABA_A_ receptors mediate rapid responses to the neurotransmitter GABA and are robust regulators of the brain and spinal cord neural networks that control locomotor behaviors, such as walking and swimming. In developing zebrafish, gross pharmacological blockade of these receptors causes hyperactive swimming, which has been embraced as an epilepsy model. Although GABA_A_ receptors are important to control locomotor behavior, the large number of subunits and homeostatic compensatory mechanisms have challenged efforts to determine subunit-selective roles. To address this issue, we mutated each of the eight zebrafish GABA_A_ α subunit genes individually and in pairs using a CRISPR-Cas9 somatic inactivation approach, then we examined the swimming behavior of the mutants at two developmental stages. We found that disrupting the expression of specific pairs of subunits resulted in different abnormalities in swimming behavior at the first development stage. Mutation of α4 and α5 selectively resulted in longer duration swimming episodes, mutations in α3 and α4 selectively caused excess, large-amplitude body flexions (C-bends), and mutation of α3 and α5 resulted in increases in both of these measures of hyperactivity. At the later stage of development, hyperactive phenotypes were nearly absent, suggesting that homeostatic compensation was able to overcome the disruption of even multiple subunits. Taken together, our results identify subunit-selective roles for GABA_A_ α3, α4, and α5 in regulating locomotion. Given that these subunits exhibit spatially restricted expression patterns, these results provide a foundation to identify neurons and GABAergic networks that control discrete aspects of locomotor behavior.

## INTRODUCTION

Neural networks in the vertebrate hindbrain and spinal cord rely upon a balance of excitatory and inhibitory neurotransmitter systems to orchestrate locomotion. Classically inhibitory, the neurotransmitter Gamma-AminoButyric Acid (GABA) is recognized as a key regulator of these circuits. GABA exerts its effects through two different classes of receptors, GABA_A_ and GABA_B_. GABA_B_ receptors are G-protein coupled receptors, while GABA_A_ receptors are ligand-gated ion channels that generate rapid responses to GABA. In mammalian systems, GABA_A_ receptors exhibit remarkable diversity, with each receptor thought to form a heteropentamer containing various combinations of 19 different subunits: α1-6, β1-3, γ1-3, δ, ε, π, θ and ρ1-3 (Sieghart and Sperk 2002; Simon et al. 2004; Chua and Chebib 2017). Each subunit is encoded by a discrete gene that is spatially and developmentally regulated to generate distinct, but sometimes overlapping, expression patterns (Laurie, Seeburg, et al. 1992; Laurie, Wisden, et al. 1992; Wisden et al. 1992). Several receptor subunits confer distinct biophysical and pharmacological properties, localize to synaptic or extrasynaptic sites, interact with specific cytoplasmic proteins, and contribute to different neuronal networks (Farrant and Nusser 2005; Jacob et al. 2008; Fritschy and Panzanelli 2014). While this incredible receptor heterogeneity is not fully understood, it could provide the opportunity to better understand, or even manipulate, distinct neuronal networks.

Several studies have used pharmacological blockade of GABA_A_ receptors to reveal the central roles that these receptors play in regulating the initiation, rhythmicity, frequency and duration of locomotor network output from the spinal cord. These studies have been performed in a variety of vertebrate systems. For example, in neonatal mice, application of GABA_A_ receptor antagonists to the spinal cord has been shown to cause inappropriate, bilateral discharges and regulate the onset and duration of rhythmic activity (Hinckley et al. 2005). In lamprey, application of GABA_A_ receptor antagonists to spinal cord networks increased the frequency of locomotor bursts and disrupted lengthwise coordination (Schmitt et al. 2004). In Xenopus tadpoles, GABA_A_ receptor mediated inhibition has been found to play both a tonic role in regulating the responsiveness to touch stimuli and a phasic role that stops swimming (Perrins et al. 2002; T. D. Lambert et al. 2004; Thomas D. Lambert et al. 2004). Although these and several other studies illustrate the importance of GABA_A_ receptors in controlling locomotor networks, antagonists that block the majority of receptor isoforms were used, therefore these studies are less informative about the roles of specific subunits.

Genetic inactivation of GABA_A_ receptor subunits in mice has had only limited success in identifying subunits required to mediate locomotor behavior. Although several of the 19 subunits exhibit robust expression in portions of the brain and spinal cord that mediate locomotion, few gene deletions have been found to cause abnormal locomotor behavior (Rudolph and Möhler 2004; Vicini and Ortinski 2004; Smith and Rudolph 2012). Ablation of some GABA_A_ receptor genes has been shown to cause changes in the expression of other subunits, which suggests that homeostatic adaptations may explain at least some of the deletions that show no or only subtle locomotor phenotypes (Peng et al. 2002; Kralic et al. 2006; Zeller et al. 2008; Panzanelli et al. 2011; Zhou et al. 2013; Fritschy and Panzanelli 2014).

Early larval-stage zebrafish, ∼2-10 days post-fertilization, are a leading model for locomotor neural network analysis, and GABA_A_ receptors regulate locomotion in this system. Bath application of GABA_A_ receptor antagonists, such as pentylenetetrazole (PTZ), induces hyperactive swimming, meaning a dramatic increase which is also recognized as an epileptic seizure model (Baraban et al. 2005; Baxendale et al. 2012; Cho et al. 2020). Zebrafish harbor an array of GABA_A_ subunits similar to mammals, with 23 identified subunits (Cocco et al. 2017; Monesson-Olson et al. 2018), however few mutants have been identified as important for locomotion. At 5 days post-fertilization (dpf), loss-of-function mutations in the broadly expressed γ2 subunit were reported to elicit hyperactive swimming that is behaviorally similar to PTZ exposure (Liao et al. 2019). At 7 dpf, loss-of-function mutations in β3, which is also widely expressed, caused subtle increases in spontaneous swimming (Yang et al. 2019). Although loss-of-function mutations in α1 were found to cause hyperactive swimming at 5 weeks post-fertilization, this behavior was not observed during larval stages (Samarut et al. 2018). Since GABA_A_ α subunits help form the GABA binding site, they are thought to be obligatory receptor components (Phulera et al. 2018; Zhu et al. 2018; Laverty et al. 2019; Masiulis et al. 2019). Thus, it is surprising that α subunits have not been identified as important for regulation of embryonic or larval locomotor behavior. It is possible that, as in mammals, homeostatic compensation is able to conceal subtype-selective roles. Disrupting multiple subunits simultaneously could evade these mechanisms and reveal α subunits that are required to regulate swimming behavior.

Here, we used an F0 CRISPR-Cas9, somatic mutation approach to screen the locomotor phenotypes of mutants in each of the eight α subunits both individually and in combination at two different developmental stages: 48 and 96 hours post fertilization (hpf). We found that disrupting select pairs of α subunits causes different types of hyperactive behavior at 48 hpf, which then decreased or was absent by 96 hpf. The absence of hyperactive behavior by 96 hpf was confirmed in F2 germline mutants and, correspondingly, electrophysiological recordings revealed brain activity indistinguishable from wild-type controls. These findings illustrate subunit selective-roles of GABA_A_ receptor α subunits in regulating locomotor behavior which, given their restricted expression patterns in larvae, serve as entry points to reveal cellular and circuit mechanisms that enable GABA to control locomotion.

## MATERIALS AND METHODS

### Zebrafish maintenance and breeding

Adult zebrafish were maintained according to standard procedures, with the zebrafish facility on a 14-hour light/10-hour dark cycle. Embryos and larvae were kept at 28.5°C in E3 media and staged according to morphological criteria (Kimmel et al. 1995; Parichy et al. 2009). All genetic manipulations and behavioral experiments were performed using a Tübingen (Tü) or Tüpfel longfin (TL) genetic background. All animal procedures for this study were approved by the University of Massachusetts Amherst or the Amherst College Institutional Animal Care and Use Committee (IACUC). Amherst College assurance number 3925-1 with the Office of Laboratory Animal Welfare.

### Guide and Cas9 RNA preparation and microinjection

Single guide RNAS were designed using the online tool, ChopChop v.2 (Labun et al. 2019) (Supplemental Table). ChopChop selects target sites for CRISPR-Cas9 using the NGG motif and ranks them based on efficiency (Montague et al. 2014). For each gene, two neighboring targets were selected with the high efficiency within exons that would disrupt all known splice-variants as assessed in ENSEMBL. CRISPR-Somatic Tissue Activity Tests were used to assess mutational efficiency for both targets of a selected gene, and the target that yielded better results was used. In some cases, neither of the initial targets was effectively mutated so two additional targets were selected.

Template DNA for gRNA synthesis was generated using the PCR based method described in (Shah et al. 2016). For *in vitro* transcription, we first generated an HPLC-purified scaffolding primer (5’-GATCCGCACCGACTCGGTCCCACTTTTTCAAGTTGATAACGGACTAGCCTTATTTTAA CTTGCTATTTCTAGCTCTAAAAC-3’), which was common to all gRNA templateless PCR reactions. Next, for each target, we synthesized a unique primer sequence that contained a 5’ T7 binding site, the 20 nucleotide specific target (Supplemental Table), and a 3’ 20 nucleotide site of scaffolding homology (5’-AATTAATACGACTCACTATA-[20 nucleotide Target Sequence]-GTTTTAGAGCTAGAAATAGC-3’). The PCR reaction contained: 0.4 units of Phusion High-Fidelity DNA Polymerase (New England Biolabs, M0530S), 13.4 μL ddH2O, 1μL target specific primer (10μM, IDT), 1 μL scaffolding primer (10μM, IDT), 4μL 5X Phusion HF, and 0.4μL dNTPs (10mM). Reactions were run in a thermocycler (BioRad) using the following conditions: 98°C for 30 seconds then 40 cycles of 98°C for 10 seconds, 60°C for 10 seconds, 72°C for 15 seconds, which was followed by 72°C for 10 min. The PCR reaction was purified using the QIAGEN MinElute kit and used as a template for *in vitro* transcription reactions. Using 0.5-1 ug of purified PCR product, gRNAs were generated using the MEGAscript T7 Transcription kit (Thermofisher), purified via lithium chloride precipitation, and verified using a TAE denaturing gel. A nanodrop spectrometer was used to determine gRNA concentrations, which were then diluted to 200ng/ul in RNase free water.

Cas9 mRNA was synthesized similar to Shah *et al*., 2016, but with the following changes. Purification of the linearized plasmid was performed using the E.Z.N.A. Cycle Pure Kit (Omega Bio-Tek), while purification of the Cas9 mRNA was by lithium chloride precipitation. Cas9 mRNA was diluted to 1200ng/ul in RNase free water.

Microinjections were performed at the 1-4 cell stage into the yolk of embryonic zebrafish. The injection cocktail contained: 2μL gRNA at 200ng/μL, 2ul Cas9 mRNA at 1,200ng/μL, 1ul stop cassette (Gagnon et al. 2014) at 10μM, 2μL 0.05% Phenol Red, 4μL RNase free water. Approximately 500 pL was injected per embryo. Mock injections, containing all cocktail ingredients except gRNAs, were not observed to have a significant effect compared to uninjected siblings, so uninjected sibling animals were used as the controls for most experiments.

### Tyrosinase Pigmenation Analysis

24 hours after injection, injected embryos were dechorionated using forceps and screened for morphological abnormalities using a dissecting scope (Zeiss). Morphologically abnormal fish were excluded from further analysis. At 48 hours and 96 hours post-fertilization (hpf), larvae were anesthetized using 0.04% MS-222 (Tricaine) and pictures were taken using a Stemi 305 (Zeiss). All images were captured with the same dimensions (1280 × 960), resolution (72 × 72), lighting conditions and exposure times. Images were analyzed using a custom Python script (source code available upon request). Briefly, a threshold value was empirically selected based on its ability to most optimally segment pigmentation from other parts of the fish and the background across all images. The number of pixels greater than this threshold were summed and reported for each image. The same threshold value was applied to all images.

### Behavioral Analysis

Behavioral analysis was performed in a double-blind fashion. At 24hpf, injected embryos had their chorions removed and were subject to morphological screening. At 48hpf, we examined escape responses to touch responses similar to as described in (Friedrich et al. 2012; McKeown et al. 2012). Briefly, light touch was applied to the head using a 3.22/0.16g of force von Frey filament. Swimming responses were captured using a high-speed digital camera (XStream 1024, IDT Vision) mounted to a 35 mm lens (Nikon) at a frame rate of 250 Hz. The head-to-tail angle for each frame of the response was measured using custom software written in MATLAB (source code available upon request). C-bends were defined as any body flexion over 110 degrees, while escape response duration was defined as beginning the frame before initial movement was observed and ending at the last frame of detected movement (McKeown et al. 2012).

### CRISPR-Somatic Tissue Activity Tests (STAT)

CRISPR-STAT analysis was performed similar to as described by Carrington et al. 2015 (Carrington et al. 2015). Primer pairs were designed to flank the targeting gRNA sites as determined in ChopChop span target sequences (Supplemental Table). Forward primers were tagged with a 6-FAM dye (Integrated DNA Technologies) which allowed for visualization in fragment analysis. Reverse primers were tagged with a PIG-tail adapter (5’-GTGTCTT-3’) to reduce stutter (Blake et al., 2015). DNA was extracted from 6 embryos using the Extract-N-Amp Tissue PCR Kit (Millipore-Sigma) and amplified using Amplitaq Gold Taq Polymerase (ThermoFisher). PCR conditions followed those suggested by the taq polymerase manufacturer. DNA was diluted 1:20 in ddH_2_0 and ran on an AB3730xl DNA Analyzer (Genewiz, South Plainfield, NJ). Results were analyzed using Geneious Software (Biomatters, Inc). Peaks were defined as signals exceeding 1,000 Relative Fluorescent Units (RFU). A gene target was considered successfully mutated if the peak 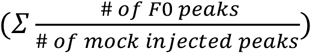 ratio was 2 or above.

### Gabra3 MUTANT LINES

To generate *gabra3* mutant lines, CRISPR-Cas9 injected animals were raised to adulthood. CRISPR-STAT analysis was used to identify mosaic animals and these animals were crossed to a wild-type strain (TLF). Two *gabra3* mutant alleles were identified in the F1 animals, a 7-base pair deletion (bp), *uma500*, and an 18-bp insertion, *uma501*.

### Local Field Potential (LFP) Recording

Local field potential (LFP) recordings were obtained from zebrafish larvae at 96 hpf using a technique similar to that in Liu and Baraban, 2019. Prior to each recording, larvae were paralyzed by immersion for 30-60 minutes in α-Bungarotoxin (125 µM in dH20Invitrogen, Waltham MA) and were subsequently embedded dorsal side up in 2% low-melting-point agarose in extracellular solution (130 mM NaCl, 10 mM HEPES, 2 mM KCl, 2 mM CaCl2, 1 mM MgCl2, pH 7.8). In some experiments, the convulsant agent PTZ (10 mM) was applied to induce ictal-like brain activity. For each recording, a glass microelectrode (6-12 MΩ) was filled with extracellular solution and inserted under visual guidance into the optic tectum. Local field potentials were recorded at 100X gain in current clamp mode using a Sutter Double IPA amplifier (Sutter Instruments, Novato CA). Voltage signals were low-pass filtered at 500 Hz-1 kHz and digitized at 5-10 kHz using SutterPatch Software (Sutter Instruments, Novato CA). Following acquisition, voltage traces were analyzed for ictal-like activity using NeuroMatic software (Rothman and Silver 2018) and the number of ictal-like events in the first 30 minutes of recording was assessed. For PTZ treated fish, one hour was allocated for wash-on and only the first 30 minutes of recording after this wash-on period was analyzed.

### Statistics and Analysis

To determine significant differences the following statistical tests and software were used. Welch’s t-test and Ordinary 1-way ANOVA were used where appropriate. When t-tests were applied, F tests were used to compare variance. When ANOVAs were applied, multiple comparison tests were used where test groups were compared against wildtype controls. A Dunnett test was used to correct against familywise error. Figures, plots and statistical testing were performed using Prism (GraphPad Software).

## RESULTS

### High-efficiency F0 somatic gene targeting using CRISPR-Cas9

To identify GABA_A_ α subunits that control larval escape behavior we sought an approach to rapidly screen through different loss-of-function mutant combinations. Injecting zebrafish embryos with a cocktail containing Cas9-encoding and guide RNAs (gRNAs) has been shown to mutate target genes with enough efficiency to cause biallelic disruption (Shah et al. 2015; Varshney et al. 2015; Wu et al. 2018). Phenotypes can often be observed in these F0 somatic mutants, which can save a great deal of time compared to analysis of F2 germline homozygous mutants. A disadvantage of this approach is that F0 somatic mutants are genetic mosaics, with different cells harboring different indels in the target gene, which may yield weaker phenotypes compared to those found in germline mutants. To confirm the efficiency of disrupting gene function using this approach, we targeted the tyrosinase (*tyr*) gene, similar to previous studies (Wu et al. 2018). Tyrosinase is essential for producing melanin and its disruption provides an easily observable loss of pigmentation. Phenylthiourea (PTU) is routinely used in zebrafish research to suppress melanin synthesis and it provides an effective method to evaluate CRISPR knockdown of *tyr*. We observed that injection of Cas9 and gRNAs targeting the *tyr* gene led to a substantial and persistent reduction in pigmentation in 48 and 96 hpf larvae, although not all melanin synthesis was eliminated compared to PTU treated controls (Supplementary Figure 1A, B). Correspondingly, PCR analysis of the somatic mutants through CRISPR-Somatic Tissue Activity Tests (STAT), indicated that a variety of mutations were induced in the *tyr* gene (Supplementary Figure 1C). Taken together, these results confirm that F0 somatic mutants provide a rapid and effective means to screen gene function.

### GABA_A_ Receptors Control Early Larval Swimming Behavior

Although PTZ is known to elicit hyperactive swimming responses in fish older than 5 dpf, its effect on earlier larval stages is less clear. We focused behavioral analysis on 48 and 96 hpf larvae since these time points present the opportunity to analyze a more nascent zebrafish nervous system and the early stages of GABA_A_ receptor control of locomotion. At 48 hpf, larvae are newly hatched and demonstrate burst swimming behavior, while 96 hpf larvae exhibit more mature swimming patterns (Brustein et al. 2003; McKeown et al. 2009; Roussel et al. 2020). Because acoustic and light responses are either absent or less robust at these early developmental stages, light touch was used to induce escape response swimming behavior. At both 48 and 96 hpf, larvae respond to light touch to the head with a well-characterized C-start, which consists of an initial C-shaped body bend to reorient the animal away from the touch stimulus, followed by lower-amplitude body undulations that propel it several body lengths away (Eaton et al. 1977; Granato et al. 1996; O’Malley et al. 1996; Eaton et al. 2001; Kohashi et al. 2012). After PTZ exposure, hyperactive behavior is observed, and two prominent aspects of swimming performance are altered similar to older fish: larvae performed longer duration escape responses and these responses are interspersed with multiple C-shaped body bends (Figure 1; Supplementary Movies 1 and 2). These results indicate that GABA_A_ receptors are essential to regulate swimming behavior during escape responses in early larval zebrafish, as has been previously shown for later stages of development.

**Figure 1.**
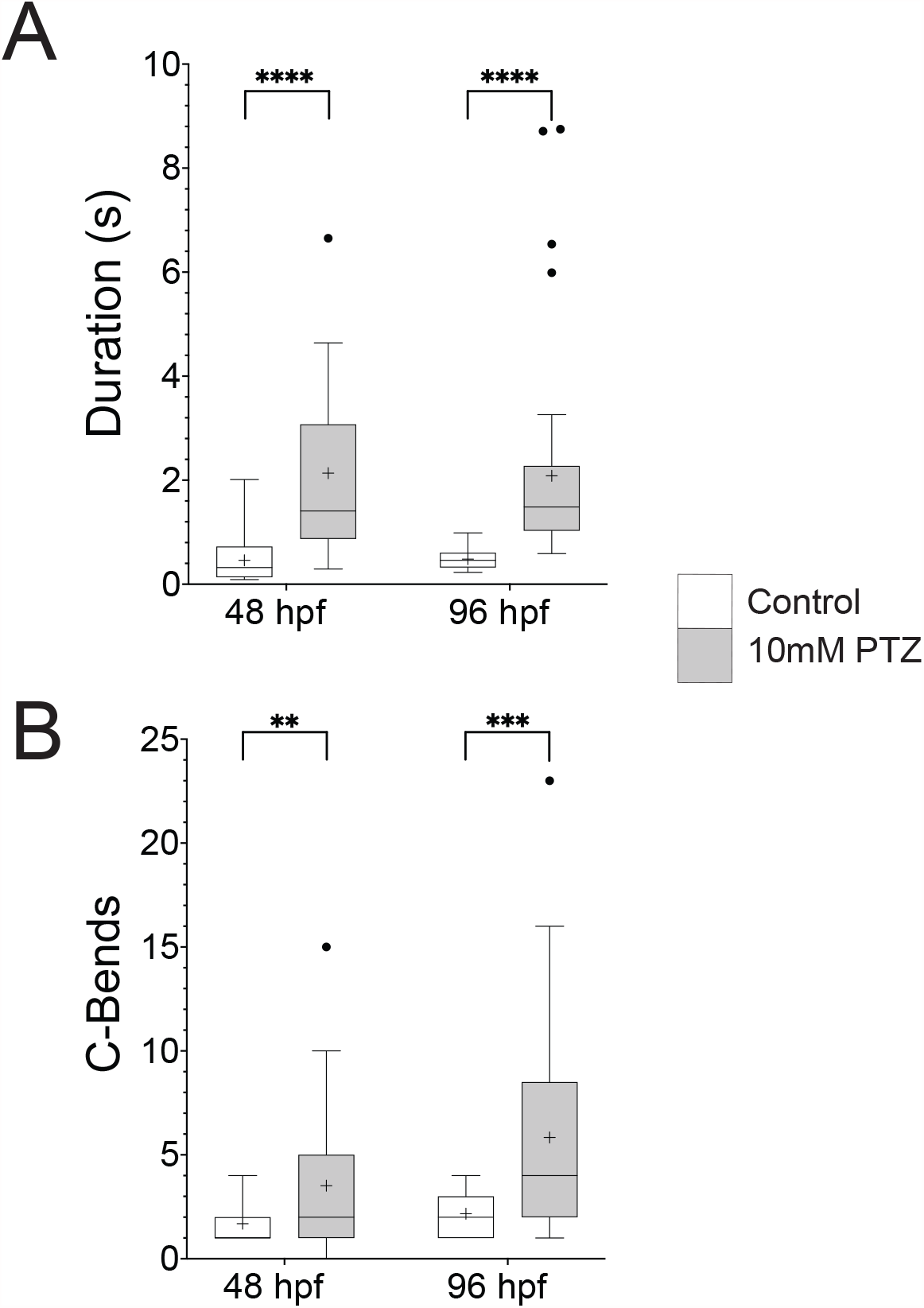
Pentylenetetrazole (PTZ) exposure induces hyperactive swimming in early larval zebrafish. PTZ is a potent GABA_A_ receptor antagonist. Bath application of 10 mM PTZ to both 48 and 96 hpf zebrafish caused an increase in (A) swim duration shown in seconds and (B) C-bends, defined as large-amplitude body bends over 110°. Box plots are representing the 25% and 75% quartiles with the median represented by a horizontal black line and the mean represented by a black plus sign within the box. Tukey’s whiskers were used. *n*=20 and 39 for wild type and PTZ treated larvae, respectively, \*\**P*<0.01, \*\*\**P*<0.001, \*\*\*\**P*<0.0001 using unpaired Welch’s t-tests.

### Mutation of pairs of α subunits induces hyperactive behavior at 48 hpf

To identify GABA_A_ α subunits that control locomotion, we generated F0 somatic mutations in each of the 8 subunits individually and in all pairwise combinations, then we analyzed touch-evoked behavior, focusing on escape response duration and body-bend amplitude (Figure 2A). Of the 36 conditions examined, no individually mutated α subunit gave rise to abnormal response durations at 48 hpf, however mutating pairs α3/α5 or α4/α5 resulted in increases in swimming times (Figure 2B). Sibling controls exhibited an average swimming duration of 0.53±0.03 seconds (n=256). In contrast, larvae with mutations in α3/α5 and α4/α5 responded with an average of 1.27± 0.33 seconds (n=18) and 1.69±0.49 seconds (n=18), respectively (Figures 2C, 2D, Supplementary Movies, 3, 4).

**Figure 2.**
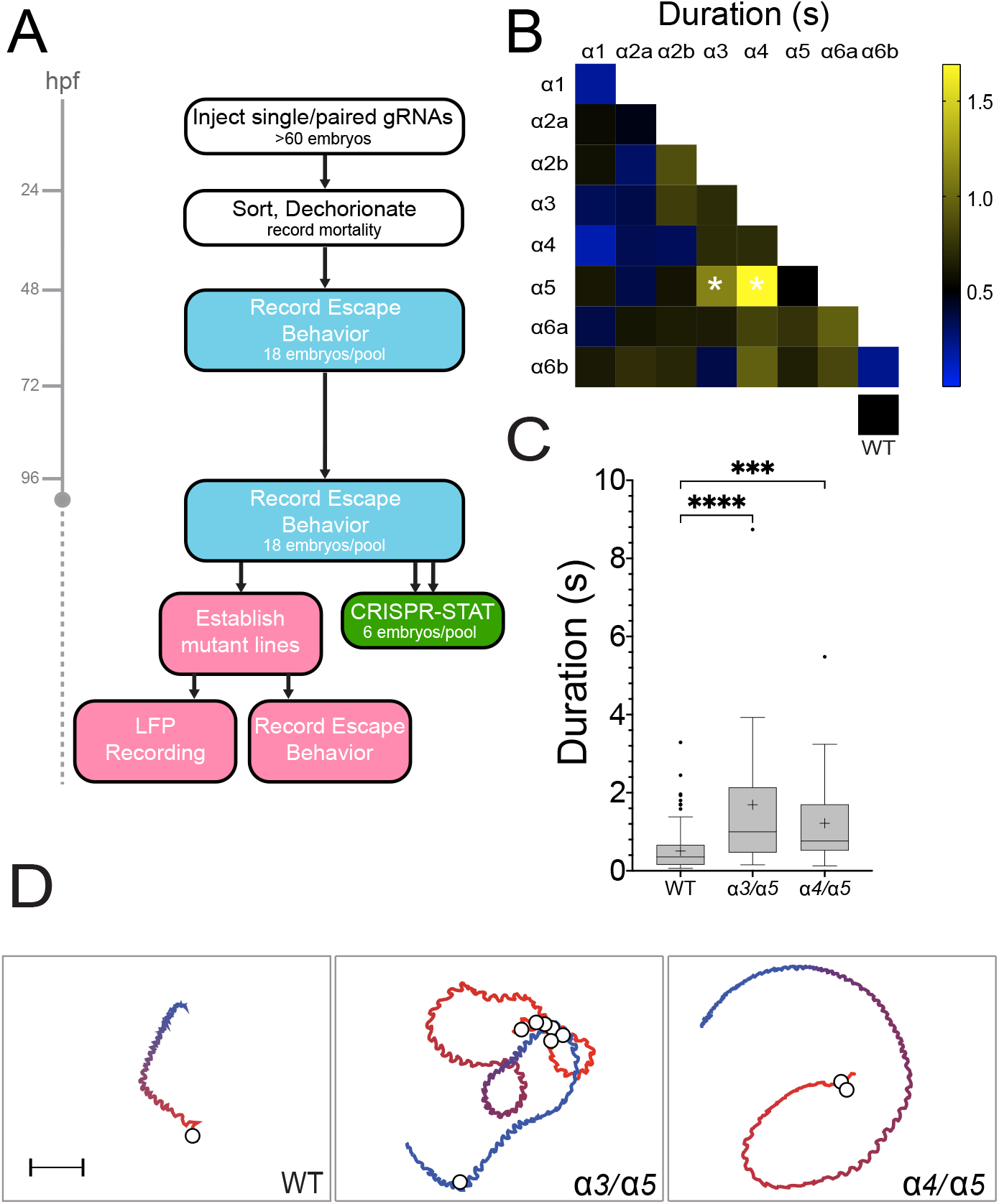
An F0 somatic mutation screen of GABA_A_ receptor α subunits identifies mutant combinations that show increased swimming durations at 48 hpf. (A) Overview of the GABA_A_R α subunit screen. Different colored boxes represent different aspects of the screen. (B) Heat matrix of the 36 single and double F0 somatic mutant conditions indicating mean startle response durations. The heatbar (*right*) indicates the average swim length. The box at lower right shows mock or uninjected controls. Ordinary one-way ANOVA revealed significant differences in swimming durations according to knock-down target, (N=995 larvae total, 612 mutants with 13-26 larvae per condition, 383 wild-type siblings; (F(36,958) = 5.316, p<0.0001). A Dunnet’s post-hoc test revealed significant pairwise differences between α3/α5 compared to wild-type and α4/α5 compared to wild-type (*white asterisks*), with those conditions exhibiting average swim durations of 1.27± 0.33 seconds (n=18) and 1.69±0.49 seconds, respectively. (C) Boxplots of α3/α5 and α4/α5 somatic mutant swimming durations show the increased swimming durations compared to controls. \*\*\**P*<0.001, \*\*\*\**P*<0.0001 using Dunnett’s multiple comparison test. (D) Traces of representative escape responses for wild-type, α3/α5 and α4/α5 somatic mutants. The color spectrum of each trace indicates the beginning (*red*) and end (*blue*) of the response, and *white circles* represent the location of C bends. The videos used to generate these traces are provided in the Supplementary Data.

Mutations in α3, α4, and α5 also caused hyperactive increases in the number of C-shaped body bends at 48 hpf. Wild-type larvae exhibited an average of 1.37 ±0.04 (n=383) C-bends per escape response, while mutations in α3/α4 and α3/α5 performed an average of 5.32±1.89 (n=18) and 3.88±1.01 (n=18) C-bends per escape response, respectively (Figure 3A, B). Notably, mutations in pair α3/α4 increased the number of C-bends per escape response without causing significantly longer swimming durations, while mutations in pair α4/α5 caused an increase in swimming durations without a significant increase in C-bends. No mutant pairs without α3 were found to increase the number of high-amplitude body bends. Similarly, no mutant pairs without α5 were found to increase swimming durations. These observations suggest that α3 might play a dominant role in controlling the number of C-bends during an escape response, while α5 predominantly regulates swimming durations.

**Figure 3.**
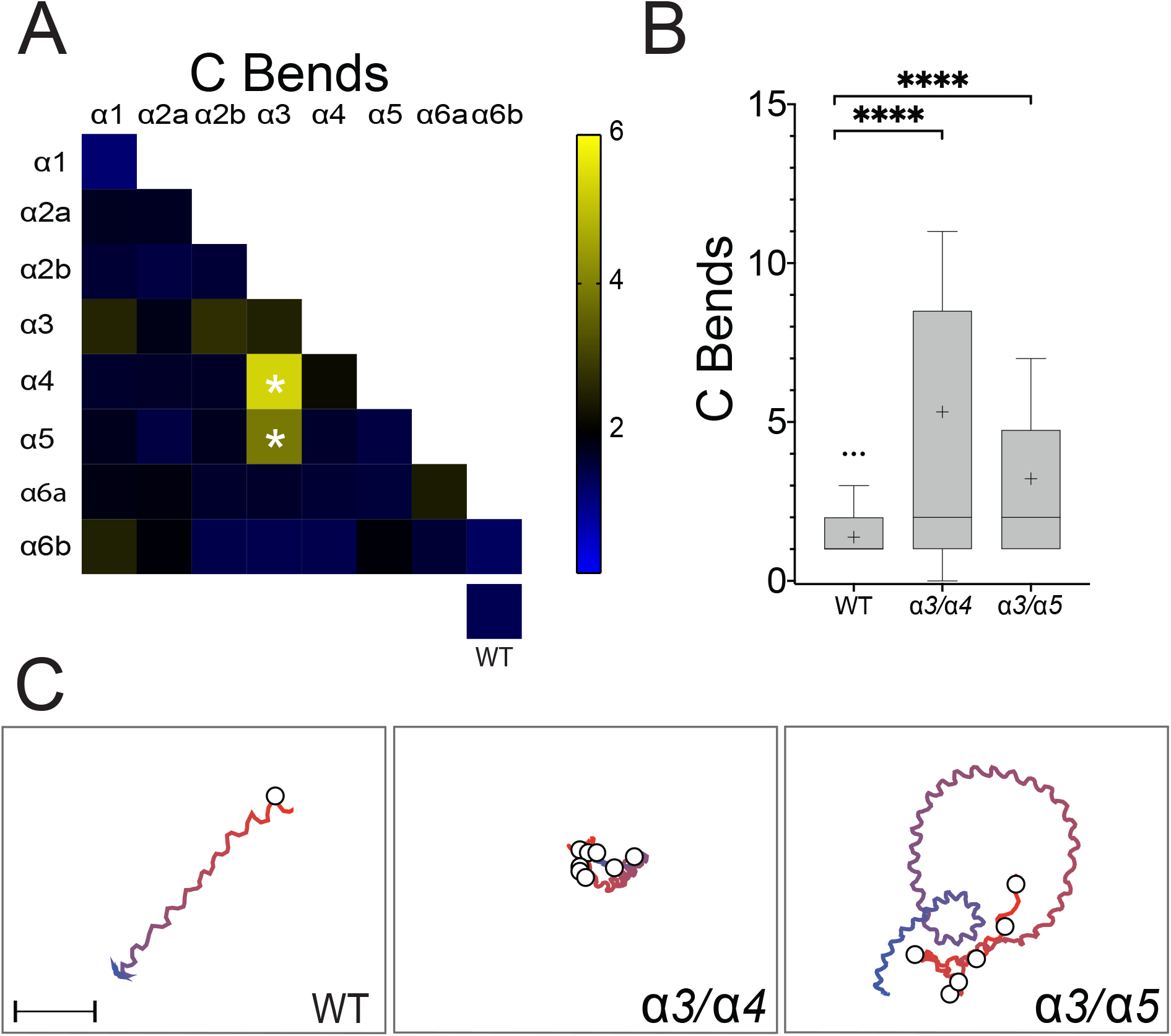
Somatic mutation of pairs α3/α4 or α3/α5 causes an increased number of large-amplitude body bends at 48 hpf. (A) Heat matrix of the α subunit single and double somatic mutant conditions indicating mean numbers of C-bends per response. The heatbar (*right*) indicates the average swim length. The box at lower right shows mock or uninjected controls. Ordinary one-way ANOVA revealed significant differences in large-amplitude body bends according to knock-down target, (F(36, 834)=4.221, p<0.0001). A Dunnett’s post-hoc test revealed significant pairwise differences between α3/α4 compared to wild-type and α3/α5 compared to wild-type (*white asterisks*), with those conditions exhibiting average C-bend per response of 5.32 and 3.88, respectively, compared to 1.37. (B) Boxplots of α3/α4 and α3/α5 somatic mutant C-bends show the increased number of large amplitude body bends per swimming episode compared to sibling controls. \*\*\**P*<0.001, \*\*\*\**P*<0.0001 using Dunnett’s multiple comparison test. (C) Traces of representative escape responses for wild-type, α3/α4 and α3/α5. The color spectrum of each trace indicates the beginning (*red*) and end (*blue*) of the response, and *white circles* represent the location of C bends. The videos used to generate these traces are provided in the Supplementary Data.

### Hyperactive phenotypes at 48 hpf are absent or attenuated by 96 hpf

The hyperactive phenotypes observed at 48 hpf were absent or greatly reduced by 96 hpf. Mutations in pair α3/α5 resulted in significantly longer duration swimming episodes at 48 hpf, however at 96 hpf the swimming behavior of these same animals was indistinguishable from controls (Figure 4A, B). Mutations in pair α4/α5 did result in longer duration swimming responses at both 48 and 96 hpf, however, at the later time point, the difference compared to controls was far less (a difference of 1.18±0.15 (n=18) seconds at 48 hpf and 0.41±0.07 (n=18) seconds at 96 hpf compared to controls). No other mutations caused increased swimming durations. Similarly, although mutations in pairs α3/α4 or α3/α5 induced excess C-bends at 48 hpf, neither these nor any other mutations were found to cause significantly greater large-amplitude body flexions at 96 hpf (Figure 4C,D).

**Figure 4.**
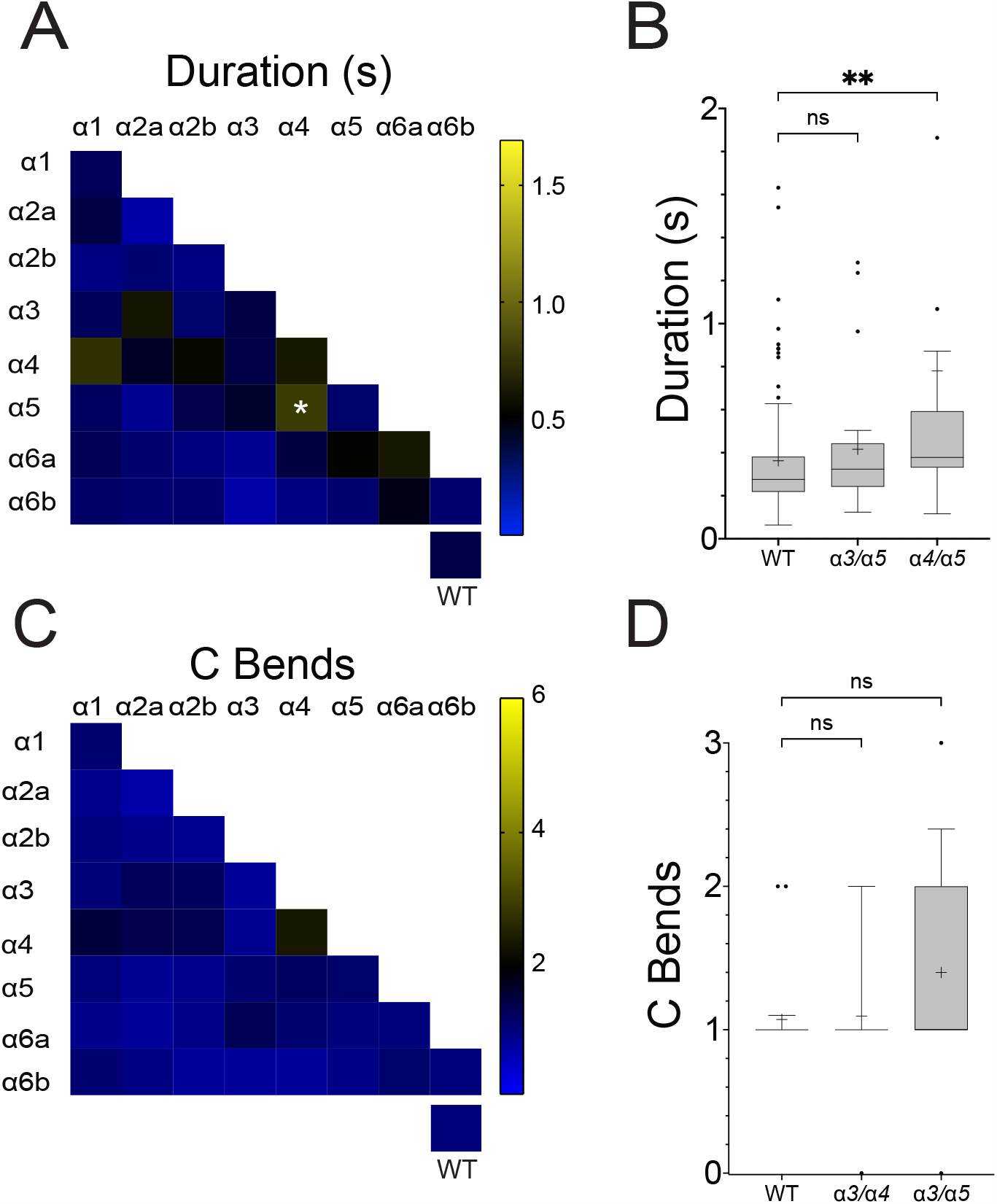
Only α4/α5 somatic mutants continue to exhibit a hyperactive phenotype at 96 hpf. (A) Heat matrix of the single and double F0 somatic mutant conditions showing evoked swimming response durations at 96 hpf. The heatbar (*right*) indicates the average swim length. The box at lower right shows mock or uninjected controls. Ordinary one-way ANOVA revealed a significant difference in swimming duration dependent upon knock-down target (F(36,754) = 2.914, p<0.0001). A Dunnet’s post-hoc test revealed only a significant pairwise difference between α4/α5 (*white asterisk*) and wild-type. (B) A boxplot of α3/α5, α4/α5 mutant pairs and controls. Both mutant conditions showed increased swimming durations at 48 hpf, however only the α4/α5 pair was statistically significant at 96 hpf. Although significant, the α4/α5 swimming duration is reduced compared to 48 hpf. \*\**P*<0.01 using Dunnett’s multiple comparison test. (C) Heat matrix of C bends at 96 hpf. No significant differences were detected. (D) Box plots show that conditions that demonstrated increased C bends at 48 hpf were not significantly elevated at 96 hpf.

### α3 F2 germline mutants confirm that hyperactive phenotypes at 48 hpf are reduced by 96 hpf

Previous studies have suggested that phenotypes observed in F0 somatic mutants can weaken during development, possibly due to the effects of mosaicism (ref). To address whether this mechanism could explain the reduction in hyperactive phenotypes at 96 hpf, we generated α3 germline mutants (Figure 5A). In line with α3 selectively controlling high-amplitude body bends but not swimming duration, α3 germline mutants demonstrated an increase in the number of C-bends without a significant increase in response times at 48 hpf (Figure 5B). The α3 germline mutant phenotype was more robust than the α3 F0 somatic phenotype, likely due to the mosaicism of somatic mutants. Similar to our observations using somatic mutants, at 96 hpf the number of C-bends were attenuated and not significantly different from sibling controls (p = 0.3879). These data indicate that the absent or weaker phenotypes observed in the F0 somatic mutants at 96 hpf are not due to mosaicism.

**Figure 5.**
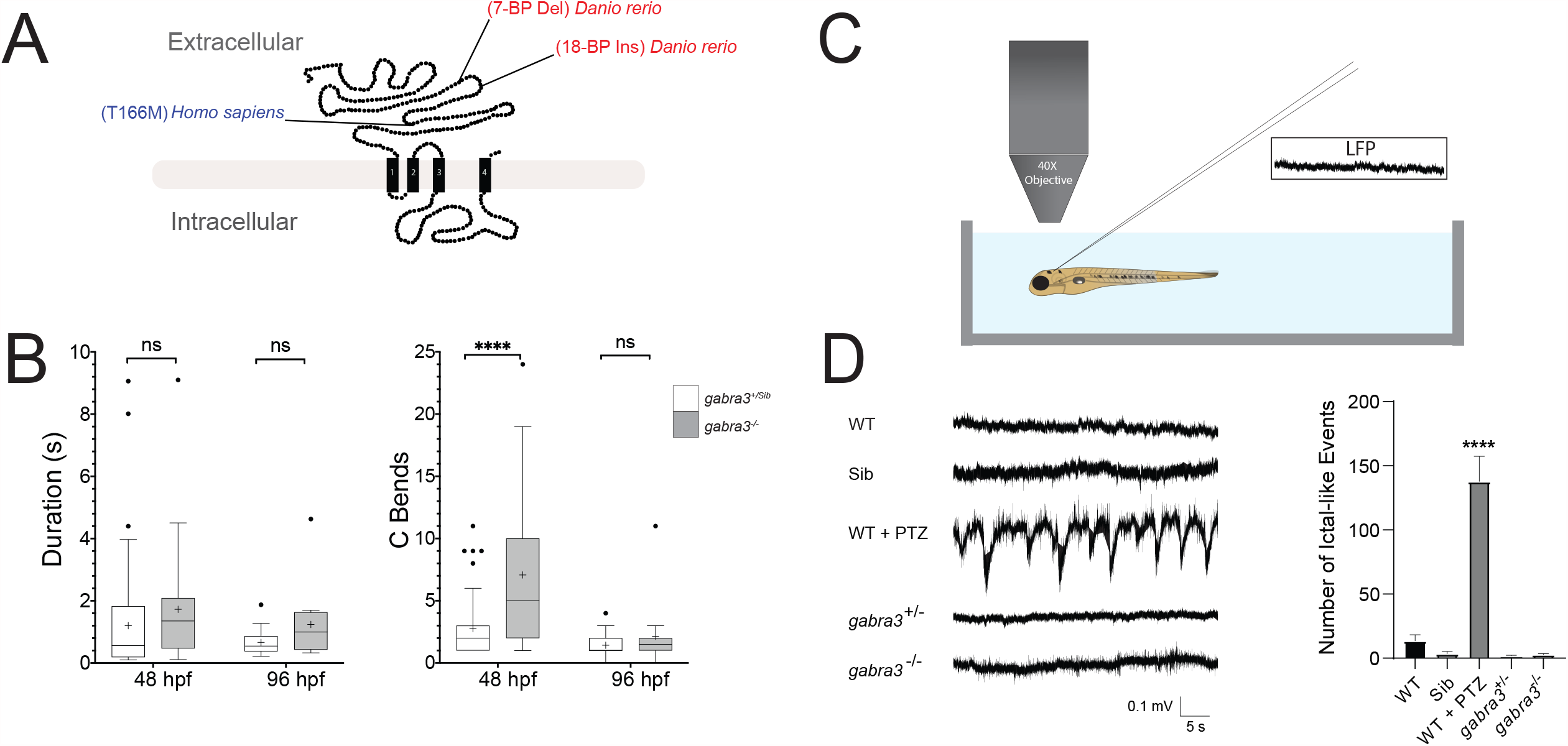
α3 F2 germline mutants confirm the α3 G0 somatic mutant phenotype. (A) A schematic of the α3 protein is shown based upon (Macdonald et al. 2010). The four transmembrane domains are indicated, along with the location of the zebrafish (*Danio rerio*) and human (*Homo sapiens*) α3 mutations. (B) Box plots for α3 trans-heterozygous mutants and siblings swimming duration in seconds (*left*) and C-bends per response (*right*) at both 48 and 96 hpf. Unpaired Student’s t-test indicated that α3 mutants exhibit significantly more C-bends at 48 hpf but not at 96 hpf ****=p<0.0001. α3 mutant swimming durations were not significantly greater than sibling controls at either time point. (C) Schematic of LFP recording setup from a 96 hpf larvae. (D) LFP traces (*left*) from wild-type (n=3), siblings (n=7), PTZ treated wild-type (n=3), α3 heterozygotes (n=6), and α3 trans-heterozygous mutants (n=3). Ictal-like activity was only detected in PTZ treated fish compared to wild-type (*right*), as revealed by ordinary one-way ANOVA with Dunnet’s post-hoc test, ****=p<0.0001.

Given that the behavior of α3 mutants was indistinguishable from wild-type controls at 96 hpf, we next asked whether local field potential recordings from the larval brain (Figure 5C) would provide a more sensitive measure of phenotype abnormalities than our behavioral assay. Consistent with previous findings (Liu and Baraban, 2019), LFP recordings over a period of 30 minutes revealed frequent large and abnormal discharges of activity at 96 hpf when PTZ was applied. However, α3 germline mutants and controls had neuronal activities that were indistinguishable from one another, consistent with their functionally normal swimming behaviors (Figure 5D).

## DISCUSSION

In this study, we performed an F0 somatic mutant screen to identify GABA_A_ α subunits that regulate hyperactive swimming during early larval stages of zebrafish development. We presented evidence that combinations of α3, α4 and α5 have selective roles in mediating different aspects of hyperactive behavior at 48 hpf. F0 somatic mutations in pairs α3/α4 significantly increased the number of high amplitude body bends, while mutations in pairs α3/α5 significantly increased swimming duration, and mutation of pairs α3/α5 caused significant increases in both parameters. We found that hyperactivity caused by somatic disruption of GABA_A_ α subunits is ostensibly reduced by 96 hpf, a result confirmed using germline α3 mutants using both behavioral and electrophysiological assays. Taken together, these data lay a foundation to investigate how GABA_A_ receptors establish and maintain control of escape behavior at neuronal and circuit levels.

### GABA_A_ Receptor Subunits Likely Control Escape Behavior Through Different Cellular Mechanisms

The GABA_A_ α subunits α3, α4, and α5 regulate escape behavior at 48 hpf, however the cellular mechanisms through which they exert their effects are not yet clear. The zebrafish hindbrain, in particular the Mauthner Cell and its homologs, play a central and well-studied role in C-starts, and these cells are regulated by GABA (Triller et al. 1997; Korn and Faber 2005; Burgess and Granato 2007; Kohashi et al. 2012; Roy and Ali 2014; Liu and Baraban 2019). α3, α4, and α5 are all expressed in discrete populations of hindbrain cells by 48 hpf, so it is possible their reduced expression dysregulates hindbrain circuits to generate hyperactive behavior (Monesson-Olson et al. 2018). Alternatively, each of these subunits is also expressed in distinct cell types in the spinal cord, so it is also possible that spinal cord GABA_A_ α subunit disruption elicits hyperactive behavior. In future studies it will be interesting to determine the relative contribution of the hindbrain versus the spinal cord in generating the abnormal behaviors caused by reduced GABA_A_ receptor function.

Whether through the hindbrain or spinal cord, α3, α4, and α5 likely control escape behavior through different neurons or subcellular mechanisms. In both the hindbrain and spinal cord at 48 hpf, α3 and α5 are expressed in overlapping domains, raising the possibility they are expressed in at least some of the same cells (Monesson-Olson et al. 2018). In both structures, α4 seems to be expressed in cells distinct from α3 and α5, therefore its effects could be mediated by different neurons.

Even when expressed in the same cells, α3, α4, and α5 are probably expressed in different subcellular domains. In mammalian neurons, GABA_A_ receptors have been shown to cluster in either synaptic or extrasynaptic domains to mediate phasic or tonic inhibition (Farrant and Nusser 2005; Jacob et al. 2008; Fritschy and Panzanelli 2014). α3-containing receptors are enriched at synapses, to mediate phasic inhibition, while α4 and α5-containing receptors are predominantly extrasynaptic, to provide tonic inhibition. Although the subcellular localizations of these subunits have not been directly investigated in zebrafish, the high degree of amino acid sequence similarity suggests that their subcellular distributions are likely conserved, which would localize α3 towards synapses while α4 and α5 would be found in mainly extrasynaptic domains. High resolution expression analysis will be required to determine if this is the case.

We did not find significant effects in response to somatic mutation of α1, α2a, α2b, α6a or α6b, however these subunits cannot be entirely ruled out from playing regulatory roles in controlling zebrafish escape behavior. F0 somatic mutations reduce, but do not eliminate expression, so it is possible that further reducing the expression of these subunits could reveal locomotor phenotypes. Additionally, our screen focused on two behavioral parameters at two developmental stages. Examining additional developmental stages, responses to other sensory stimuli, or other parameters, such as body bend frequency or frequency variability, could reveal roles for these subunits in controlling escape behavior.

### Zebrafish Exhibit Robust Homeostatic Mechanisms Across Development

The hyperactive phenotypes observed at 48 hpf were all absent or greatly reduced by 96 hpf. This result was surprising since PTZ readily elicits hyperactive behavior at all time points after 48 hpf, demonstrating that GABA_A_ receptors play critical roles in regulating locomotion across a wide variety of developmental stages. A likely explanation is that zebrafish employ robust homeostatic compensation. The mechanisms that underlie this compensation are probably multifaceted. The teleost lineage has undergone genome duplication such that there are many duplicated genes in zebrafish, including several GABA_A_ receptor subunits (Amores et al. 1998; Postlethwait et al. 1998; Monesson-Olson et al. 2018). The expression of homologous genes or simply genes with similar sequence motifs can be recruited through transcriptional adaptation, which is thought to be triggered by nonsense mediated decay (El-Brolosy et al. 2019). Mutations that cause frame-shifts and premature stop codons can cause nonsense mediated decay, therefore many of the mutations generated in this study almost certainly induced transcriptional adaptation. Pairs of α subunits were mutated to uncover adaptations within the α subunit subfamily, however transcriptional responses could involve other GABA_A_ receptor subunits. At the network level, neurons could switch the neurotransmitter they release or entire circuits could be reconfigured to maintain excitation-inhibition balance as has been observed in developing frogs and some α subunit knock-out mice, respectively (Schneider Gasser et al. 2007; Panzanelli et al. 2011). Our results here identify a narrow window between 48 and 96 hpf across which robust adaptations occur in developing zebrafish. The short time period, relatively small nervous system, ex-utero development, genetic resources, and high-resolution brain and spinal cord atlases make larval zebrafish an outstanding system to further investigate the homeostatic mechanisms activated by GABA_A_ receptor mutation.

### Zebrafish α3 Mutants as a Possible Epilepsy Model

In addition to their roles in modulating locomotion, GABA_A_ receptors are widely viewed as central factors in the development, progression, and treatment of epilepsy syndromes (Olsen and Avoli 1997; Treiman 2001; Cherubini 2012; Walker and Kullmann 2012). Abnormalities in GABA_A_ receptor inhibition are observed in genetic and acquired epilepsies, drugs that block these receptors, like PTZ, cause seizures, and drugs that enhance GABA_A_ receptor inhibition are potent anticonvulsants. In zebrafish, PTZ application is an established seizure model, and loss-of-function mutations in γ2 and β3 cause larval hyperactive behavior and/or neuronal activity that model the epilepsies caused by mutations in their human orthologs (Baraban et al. 2005; Baxendale et al. 2012; Liao et al. 2019; Yang et al. 2019; Cho et al. 2020). Similarly, mutations in zebrafish α1 cause light-triggered, hyperactive behavior in juvenile fish (older than ∼5 weeks) that seems to model epilepsy caused by reduced α1 function (Samarut et al. 2018). Here, we showed that mutation of α3 causes hyperactive behavior in larval zebrafish. Human loss-of-function variants in α3 cause a rare, severe epileptic encephalopathy, raising the possibility that zebrafish α3 mutants at least partially model this disorder (Niturad et al. 2017). It is not yet clear whether the zebrafish hyperactive swimming observed at 48 hpf is due to brain or spinal cord mechanisms but, regardless, it is relatively simple to screen the phenotype caused by α3 disruption. Given the proven effectiveness of larval zebrafish for high-throughput small molecule screens, α3 mutants could be a new and useful resource to identify novel anti-epileptic drugs (Griffin et al. 2017; Lam and Peterson 2019; Griffin et al. 2020; Patton et al. 2021 Jun 11).

## ACKNOWLEDGEMENTS

The authors thank Abhay Mittal for developing kinematic analysis software; Saige Calkins, Caroline Martin, and Oshiomah Oyageshio for excellent fish care; and Marie Abate, Sean Doherty, Ana Dolan, and Chinemerem Nwokemodo-Ihejirika for technical assistance. We also thank the rest of the members of the Downes and Trapani labs, and the entire University of Massachusetts zebrafish community for thoughtful discussion.

## FIGURE LEGENDS

**Supplementary Figure 1. CRISPR-Cas 9 targeting of *tyr* confirms high-efficiency gene targeting**. (A) Wild-type uninjected (*left*) and fish in which tyr gRNA and Cas9 encoding RNA were injected (*right*) at 48 (*top*) and 96 hpf (*bottom*). The fish in which the *tyr* gene was targeted show reductions in melanophores. (B) Bar graph showing a pixel density analysis of uninjected (n=21), *tyr* CRISPR*-*injected (n=36), or PTU-treated larvae (n=20). *tyr* CRISPR-injected larvae show reduced pixel density at both developmental ages, indicating effective gene knock-down. Each group measured against pixels detected above an arbitrary threshold determined *a priori*. Ordinary One-Way ANOVA was used ****=p<0.0001. (C) Fragment Analysis of WT (*top*) or *tyr* injected embryos (*bottom*). While PCR analysis of wild-type larvae reveals a single peak of ∼278 base pairs, PCR analysis across this same region of tyr-CRISPR-injected larvae shows several peaks, indicating the presence of indels in the target region.

**Supplementary Movie 1. Representative movie of a wild-type larva touch response at 48 hpf**. The video was recorded at 250 frames/second (1.18 seconds in length) and was used to generate the trajectory trace shown in Figure 2D. Wild-type larvae respond to touch by performing a C-bend followed by rhythmic, smaller amplitude body bends to propel the animal away from the touch stimulus.

**Supplementary Movie 2. Representative movie of an α3/α5 G0 somatic mutant at 48 hpf**. The video was recorded at 250 frames/second (5.48 seconds in length) and was used to generate the trajectory trace shown in Figure 2D. α3/α5 mutants exhibit both increased swimming durations and C-bends per response.

**Supplementary Movie 3. Representative movie of an α4/α5 G0 somatic mutant at 48 hpf**. The video was recorded at 250 frames/second (3.93 seconds in length) and was used to generate the trajectory trace shown in Figure 2D. α4/α5 mutants exhibit increased swimming durations without a statistically significant increase in the number of C-bends.

**Supplementary Movie 4. Representative movie of an α3/α4 G0 somatic mutant at 48 hpf**. The video was recorded at 250 frames/second (0.99 seconds in length) and was used to generate the trajectory trace shown in Figure 3C. α3/α4 mutants exhibit increased C-bends without a statistically significant increase in length of response durations.

