## Supplementary figures and images for "GABA_A_ α Subunit Control of Hyperactive Behavior in Developing Zebrafish"

### Supplementary Figure

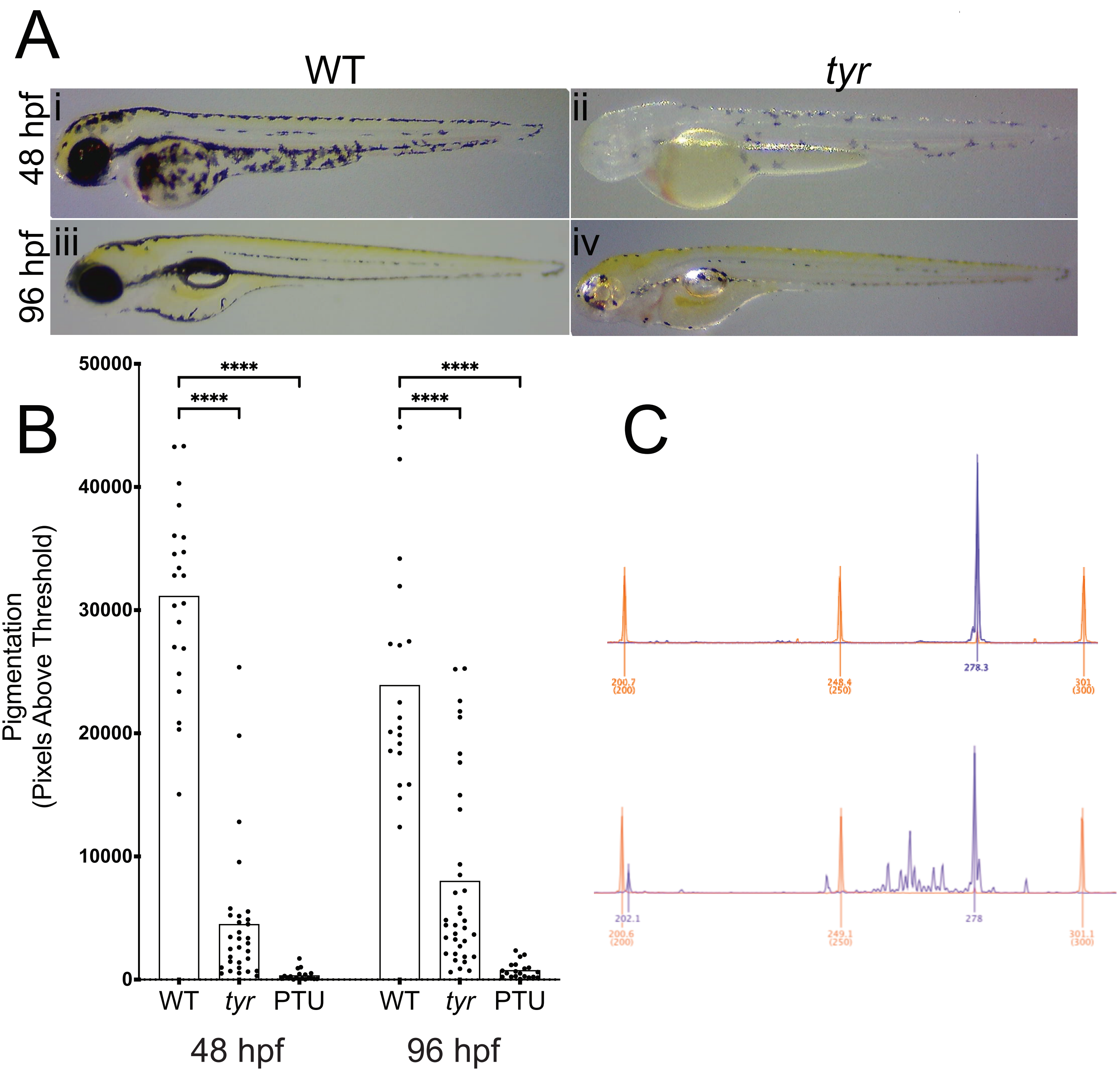
