## Supplementary Table for "GABA_A_ α Subunit Control of Hyperactive Behavior in Developing Zebrafish"

| Gene | Transcript | Exon | Target 1 (5'-3') | Target 2 (5'-3') | Flanking Forward (5'-3') | Flanking Reverse (5'-3') |
|---|---|---|---|---|---|---|
| *gabra1* | ENSDART00 000166148.2 | 5 | GCCGTTGTGGAA GAACGTGT | GAGGTTGTTGAGACG GAGCA | GAATTGTCTGTGACA AGGCTCA | ATCAAATCCAATTCT GTCCCAG |
| *gabra2a* | ENSDART00 000130007.3 | 6 | GCTCAACAGAGA GTCAGTTC | AGAGACTTGAGATAA GATGA | TCATGTTTCTCTCCT GCTGTGT | AAAGGATGTTTTATA GCCAGCG |
| *gabra2b* | ENSDART00 000192863.1 | 3 | GTAGATGTCGGT CTTCACTG | GTCTGAAACAGGGC CGAAAC | TCCTATTGAGGGATG AGCAAAT | TCTTTTTGTCTGGGT AGGTGGT |
| *gabra3* | ENSDARG00 000090883.5 | 3 | GGTATGGCTTCA GCGCTTGG | GTTGTGGAAGAAGGT ATCCG | GGACGAGAGGTTGA AGTTTGAC | TTGCATGTTTTGTAT CTCGCTT |
| *gabra4* | ENSDART00 000014454.8 | 2 | TGTCGTATCCAT CCAGGAGT | TGGCACTTACCCCCA AATCC | GCTTCAGTTTGCTCT GTGTTGT | CACTTAGTAAACAG CGTGCGAC |
| *gabra5* | ENSDARG00 000070730 | 4 | TCCCTGTAGTTG ACTGGGAG | ATGATAACAGACTTC GACCT | AACGCCTAATGGAC GAGAGATA | ACTGAAGCACGATC TTGACAAA |
| *gabra6a* | ENSDART00 000054982.7 | 2 | CTTCATTTCTCTC TGCTTAGCG | AGACTGCAAAGGTTA ACAACAGG | AGGGCTATGACAATC GACTA | GATTCTAGACGGAC TTCTTG |
| *gabra6b* | ENSDARG00 000058736 | 4 | GGTCAGCAAGAT CTGGACGC | AGATGACCGGCTGAA ATTTG | TACACGATGGATGTG TTTTTCC | GTCCATTTTTGACTT GGAGAGC |
| *tyr* | ENSDARG00 000039077 | 1 | CCCCAGAAGTCC TCCAGTCC (Jao Et Al. 2013) | N/A | CAGCTCTTCAGCTC GTCTCTC | AGCGATGGCCTTTA GTGTTTTA |
